# Femoral bone growth predictions based on personalized multi-scale simulations: Validation and sensitivity analysis of a mechanobiological model

**DOI:** 10.1101/2024.12.09.627496

**Authors:** Willi Koller, Martin Svehlik, Elias Wallnöfer, Andreas Kranzl, Gabriel Mindler, Arnold Baca, Hans Kainz

## Abstract

Musculoskeletal function is pivotal to long-term health. However, various patient groups develop torsional deformities, leading to clinical, functional problems. Understanding the interplay between movement pattern, bone loading and growth is crucial for improving the functional mobility of these patients and preserving long-term health.

Multi-scale simulations in combination with a mechanobiological bone growth model have been used to estimate bone loads and predict femoral growth trends based on cross-sectional data. The lack of longitudinal data in previous studies hindered refinements of the mechanobiological model and validation of subject-specific growth predictions, thereby limiting clinical applications.

This study aimed to validate the growth predictions using magnetic resonance images and motion capture data - collected longitudinally - from ten growing children. Additionally, a sensitivity analysis was conducted to refine model parameters.

A linear regression model based on physical activity information, anthropometric data, and predictions from the refined mechanobiological model explained 70% of femoral anteversion development. Notably, the direction of femoral development was accurately predicted in 18 out of 20 femurs, suggesting that growth predictions could help to revolutionize treatment strategies for torsional deformities.

**Statements and Declarations:** The authors have no relevant financial or non-financial interests to disclose.

## 1. Introduction

Bony deformities, particularly at the femur, are commonly observed in children with and without neurological disorders. Femoral deformities are often torsional, characterized by a misalignment of the femoral neck with the knee axis in the transverse plane. This torsion is quantified by the anteversion angle (AVA). In typically developing children, the AVA decreases during skeletal growth from approximately 40° to 20° (Bobroff et al., 1999; Fabry et al., 1973). Therefore, normative values vary with age and the distinction between healthy and pathological development is challenging. An increased, but also a decreased AVA can lead to gait abnormalities (e.g., increased internal hip rotation, in-toeing foot progression angle) and subsequently altered joint loading, as well as further progressive problems such as pain and osteoarthritis (Andriacchi et al., 2004; Bruderer-Hofstetter et al., 2015; Hudson, 2016; Sharma, 2001; Wheatley et al., 2023). While the state-of-the-art clinical intervention to correct an existing pathological AVA is a de-rotation osteotomy (Buly et al., 2018; Dreher et al., 2012), proactive non-invasive intervention strategies might be possible if pathological AVA development could be identified before its occurrence. This would allow the development of innovative intervention strategies such as gait-retraining (Kainz et al., 2024b; Uhlrich et al., 2022) to normalize bone loading in a non-invasive way and ensure typical bone growth.

With the introduction of Wolff’s law in the 19th century, it has long been understood that bone growth is influenced by mechanical loads (Wolff, 1892). The Utah paradigm of bone physiology refined Wolff’s law, recognizing that biological factors determine skeletal health and growth, but mechanical loads influence these factors and are crucial for bone growth (Frost, 2001). Shear and compressive stresses within biological tissues are key factors influencing the growth of tissues such as bone and cartilage (Carter and Beaupré, 2000; Carter and Wong, 1988; Pauwels, 1980). While many simulation studies investigating bone growth patterns focused on predicting trabecular remodelling at the micro-scale, few studies simulated bone growth on the macro-scale experienced during skeletal growth (Adachi et al., 2001; Buccino et al., 2022; Huiskes et al., 2000; Kainz et al., 2020; Yadav et al., 2021).

The shape of femur, in particular the AVA, affects the paths of muscles and therefore femoral loads (Kainz et al., 2023; Wheatley et al., 2023).

Furthermore, numerous studies have shown that the morphology of the lower limb bones can alter a person’s walking pattern (Alexander et al., 2019; Bruderer-Hofstetter et al., 2015; Carriero et al., 2009; Hudson, 2016; Mindler et al., 2021). The walking pattern has a big impact on bone loads (Carriero et al., 2014; Koller et al., 2023a), which in turn directs further bone development (Carter et al., 1996; Carter and Beaupré, 2000; Carter and Wong, 1988; Wolff, 1892). This illustrates the intricate interdependence between an individual’s musculoskeletal morphology, gait pattern, mechanical loading on biological structures, and tissue growth (Figure 1).

**Figure 1:**
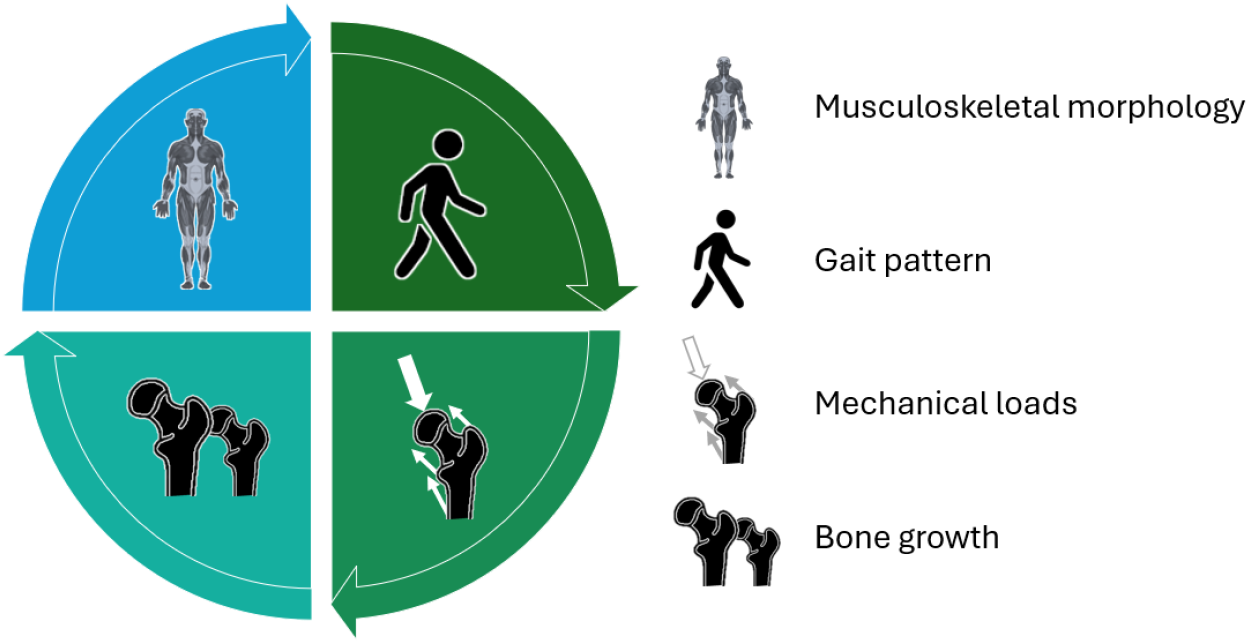
Schematic illustration of complex interplay between skeletal morphology, gait pattern, mechanical loading on bones and bone growth

Multi-scale simulations in combination with a mechanobiological bone growth model have been used to estimate bone loads and predict femoral growth trends based on cross-sectional data (Carriero et al., 2011; Kainz et al., 2020; Koller et al., 2024, 2023b; Shefelbine and Carter, 2004; Yadav et al., 2021, 2017, 2016). Within this workflow, the loadings on the participant’s femur during walking are derived from musculoskeletal (MSK) simulations based on subject-specific models and 3D gait analysis data. Subsequently, the muscle and joint contact forces from the musculoskeletal simulations are applied to a finite element (FE) model of the femur created from magnetic resonance images (MRIs) to estimate the stresses within the growth plate. Based on these stresses, a growth rate and growth direction are calculated for each element within the growth plate. Finally, growth of the femur is simulated in a second FE analysis and the development of certain angles (e.g. change in AVA) can be quantified by comparing the final with the baseline geometry (Figure 2). Previous studies using this workflow to calculate growth plate stresses and predict femoral growth revealed plausible growth patterns for healthy and pathological populations (Carriero et al., 2011; Kainz et al., 2020; Koller et al., 2024, 2023b; Shefelbine and Carter, 2004; Yadav et al., 2021, 2017, 2016). However, this workflow includes many model parameters that are based on assumptions. The lack of longitudinal data in previous studies hindered refinements of the mechanobiological model and validation of subject-specific growth predictions, thereby limiting clinical applications. The aim of this study was to close this research gap by collecting a unique longitudinal dataset, comprising 3D gait analysis data and MRIs at two time points, to experimentally investigate the subject-specific growth predictions. Furthermore, a sensitivity analysis was conducted to refine the model parameters. Multi-scale simulations were performed with two approaches to solve the muscle redundancy problem when estimating muscle forces, various parameter combinations to calculate the growth rate based on the stresses within the growth plate, and three different methods to model the growth direction. Our results demonstrate the feasibility of the refined mechanobiological model to predict femoral growth trends and show that it outperforms predictions based solely on anthropometric and physical activity data.

**Figure 2:**
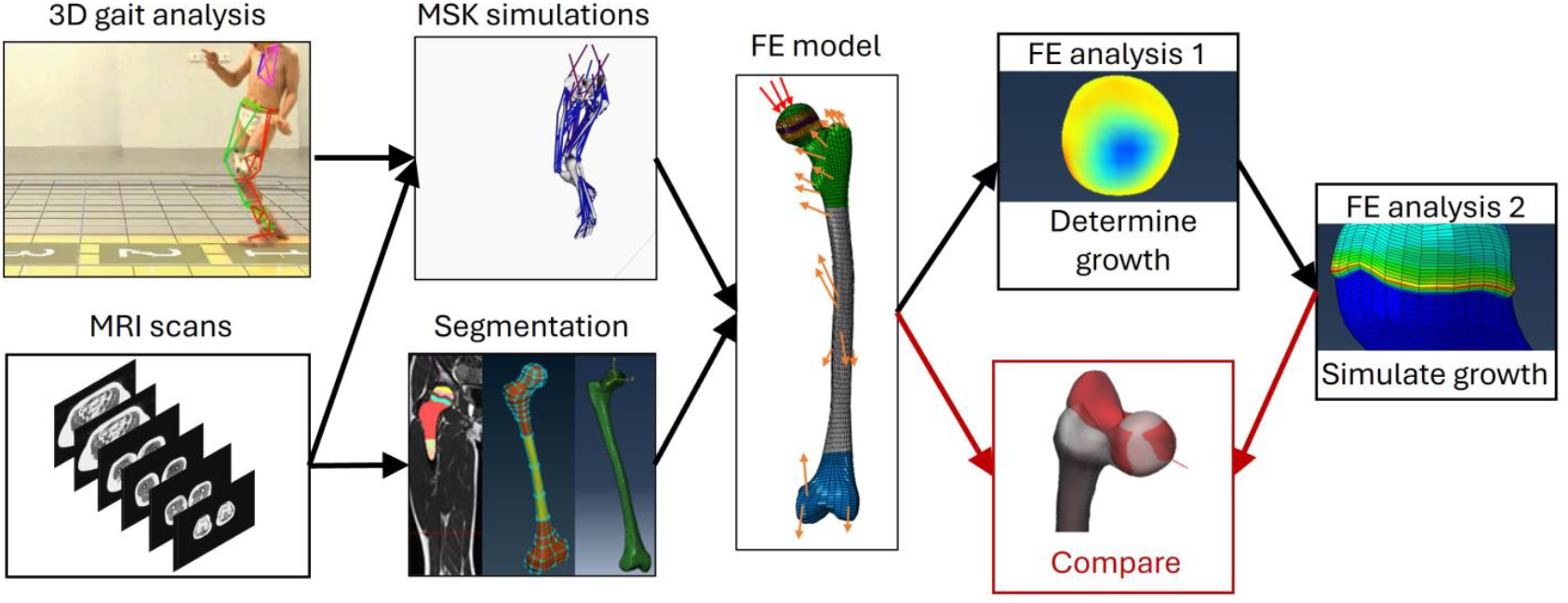
Schematic illustration of the mechanobiological multi-scale workflow to predict femoral bone growth. MSK = musculoskeletal, FE = finite element, MRI = magnetic resoncance image.

## 2. Methods

### 2.1. Data collection

Three-dimensional gait analysis data including marker trajectories, ground reaction forces and electromyography (EMG) of selected lower limb muscles as well as MRIs of the femurs of ten healthy children were collected on two occasions approximately two years apart (Table 1). Marker trajectories were collected with 200 Hz using a 12-camera motion capture system (Vicon Motion Systems, Oxford, UK). The used marker set was based on the Plug-in Gait marker set (Kadaba et al., 1990) with additional clusters of three markers on each thigh and shank segment and an additional marker at the 5^th^ metatarsal head of each foot. Ground reaction forces were simultaneously recorded with 2000 Hz with multiple force plates (Kistler, Winterthur, Switzerland). At the same time, EMG was collected with 2000 Hz of 7 lower limb muscles on each leg (tibialis anterior, gastrocnemius medialis, soleus, peroneus longus, vastus lateralis, biceps femoris, gluteus medius) with wireless surface electrodes (Cometa srl, Milan, Italy). Participants were asked to walk with a self-selected walking speed and several gait trials were recorded.

**Table 1:** Participant characteristics at first and second data collection session and the changes between session given as mean ± standard deviation and the range in brackets. BMI=Body mass index, AVA=Anteversion angle,NSA=Neck-Shaft-Angle.

| | First session | Second session | $\Delta$ between sessions |
| --- | --- | --- | --- |
| Age [years] | 9.9±0.9 (8.4 – 11.3) | 11.9 ± 0.8 (10.3 – 13) | 2.0±0.1 (1.8 – 2.1) |
| Height [cm] | 143.3 ± 7.3 (130 – 154) | 156.8 ± 8.7 (142 – 168) | 13.5±3.3 (9 - 20) |
| Weight [cm] | 35.1 ± 7.4 (29 – 50.4) | 43.3 ± 9.1 (34.2 – 62.3) | 8.2±3.3 (4.9 – 13) |
| BMI | 17 ± 2.7 (14 – 22.4) | 17.6 ± 3.1 (14.8 – 23.2) | 0.6 ± 0.9 (-0.8 – 2.1) |
| AVA [°] | 34.7 ± 8.9 (16.4 – 49.3) | 33.4 ± 9.5 (14.8 – 48) | -1.3 ± 5.8 (-13.1 – 11.8) |
| NSA [°] | 127 ± 4.2 (120.5 – 134.6) | 130.3 ± 3.5 (125.8 – 138) | 3.3 ± 3.4 (-5.1 – 8.9) |

MRIs of each femur were collected using a 3T magnetic resonance scanner (MAGNETOM Vida, Siemens, Berlin/Munich, Germany) with a T1 vibe sequence with voxel sizes between 0.8×0.8×0.7mm and 1.1×1.1×1.1mm.

Between both sessions, participants were asked to carry an activity monitoring device (Realalt 3DTriSport, London, UK) during a period of seven consecutive days to estimate their daily step count. At the second data collection session (S2), participants were requested to fill out a questionnaire about their sport and spare time activities to gain further information about their activity level. Ethics approval was obtained from the corresponding local ethics committee (University of Vienna, reference number 00578).

### 2.2. MRI measurements and segmentation of femurs

The femoral AVA in the plane perpendicular to the shaft axis and the neck-shaft-angle (NSA) were calculated based on six anatomical points of each femur defining the neck, shaft and knee axis selected in an oblique slice passing the femoral neck and in transverse slices (Sangeux et al., 2015) for both sessions. Due to uncertainties in this very common measurement methods (Kaiser et al., 2016; Sangeux et al., 2015) and to increase our confidence in the obtained AVA, the measurements were initially performed by one researcher with four years of experience in bone segmentation and anatomical feature quantification. The selected points were reviewed following the two-person verification principle by the initial researcher and a second researcher with one year of experience in bone segmentation and anatomical feature quantification. In case of disagreement, points were repositioned with consent of both researchers. The second measurement was used for further analysis and the maximum difference of the change of AVA between the two measurements was used as a measurement uncertainty (Supplementary material, Figure S1). A total of 100 imputations of the measured development of AVA were created by adding a random value within this measurement uncertainty range leading to an evenly spaced distribution. These imputations were used for subsequent analysis. Furthermore, the intercondylar distance, i.e. distance between the medial and lateral condyles of the femur, the location of the hip and knee joint centers were quantified and selected from the MRIs of the first data collection session (S1). These measurements were subsequently used to personalize musculoskeletal models which were used to estimate the loading on the femurs.

Each femur was segmented using 3D Slicer 5.2.2 (Fedorov et al., 2012) and divided into six parts – the proximal and distal trabecular bone, the proximal and distal growth plate, the cortical bone of the shaft and the bone marrow – similar to previous studies (Carriero et al., 2011; Kainz et al., 2020; Koller et al., 2023b, 2024; Yadav et al., 2016). The GP-Tool (Koller et al., 2023b) was used to identify bony landmarks (Modenese and Renault, 2021), transform the femur into the OpenSim coordinate system and create a hexahedral mesh with elements aligned with the growth plate in ten separate layers. Element size was set to 1.5 mm and 3 layers were defined as transition zones between trabecular bone and the growth plate. Linear elastic materials were assigned to the different parts of the femur with Young’s modulus and Poisson’s ratio equal to previous studies (Koller et al., 2024, 2023b).

### 2.3. Quantification of loading on bones

MSK simulations based on subject-specific MRI-informed models were performed with OpenSim (Delp et al., 2007). A modified Rajagopal model (Rajagopal et al., 2016) which included the more complex knee joint of the Lerner model (Lerner et al., 2015) was used as a base model (Kaneda et al., 2023). This model includes three rotational degrees of freedom at the hip and two rotational degrees of freedom at the knee and ankle joints. The metatarsophalangeal joint was locked. The TorsionTool (Kainz et al., 2024b; Veerkamp et al., 2021) was used to modify the femoral geometry to match each child’s NSA and AVA. These modifications alter attachment and via points of muscles acting on or passing the femur and therefore also the lever arms of muscles across joints. To fit the models to the participants’ anthropometry, the pelvis and femur segments were scaled using the hip and knee joint measurements obtained from the MRIs. Other segments of the model were scaled using the location of surface markers (Kainz et al., 2017).

Inverse kinematics were used to calculate joint angles of corresponding gait trials for each participant. Markers on the knee and ankle were neglected during inverse kinematics, instead clusters of three markers were used to track the motion of the thigh and shank segments. All markers were weighted equally. Maximum marker errors and root-mean-square errors were accepted if less than 4 cm and 2 cm, respectively, as suggested by OpenSim’s best practice recommendations (Hicks et al., 2015).

Since altering muscle attachment and via points on the femur can cause irregularities in muscle moment arms, the moment arms of all muscles acting on and spanning the femur were checked for smoothness during the motions derived from inverse kinematics. Discontinuities were observed for the anterior compartment of the musculus gluteus maximus (glmax1) in some participants and were corrected by stepwise reduction of the diameter of corresponding WrapObjects and visual inspection to ensure valid muscle paths. Subsequently, the MuscleParamOptimizer (Modenese et al., 2016) was used to map the muscle characteristics from the generic model to the personalized models. Furthermore, an actuator with an optimal force of 100 N was used to represent ligaments that passively generate the knee adduction moment.

Two approaches were used to solve the muscle redundancy problem: 1) Static optimization (SO) which minimized the sum of squared muscle activations and 2) an EMG-informed approach which minimized effort and tracking errors between EMG signals and muscle excitation patterns with OpenSim Moco (Dembia et al., 2020) with weights of 1 and 5, respectively. Subsequently, contact forces acting on the hip and knee joints were estimated (Steele et al., 2012). Additional analyses were performed to identify muscle attachments on the femur and obtain the effective directions of muscle forces (van Arkel et al., 2013). For each femur, the mean waveform of the resultant hip joint contact force (HJCF) from all trials was calculated and the trial with the lowest root mean square difference to the mean waveform was selected as a representative step and chosen for further analysis.

### 2.4. Finite element simulations

The GP-Tool (Koller et al., 2023b) was used to create a subject-specific FE model based on the participant’s femoral geometry (overall femoral shape, location and orientation of growth plate) and loading condition. Similar to previous studies (Kainz et al., 2020; Koller et al., 2023b; Yadav et al., 2016) nine load instances were selected based on the HJCF peaks and the valley in-between during the stance phase. The muscle and joint forces at these nine load instances were used as loading conditions for FE simulations. The HJCF was applied as nodal forces distributed to the closest 100 surface nodes (approximately 2.25 cm^2^) in the direction of the corresponding force orientation. Muscle forces were applied as nodal forces at the identified locations and directions. The nodes of the femoral epicondyles were constrained in all directions as boundary conditions. FEBio (Maas et al., 2017, 2012) was used to estimate stresses within the growth plate for these loading conditions.

Based on a weighted linear combination of shear and hydrostatic stresses, a value representing the growth rate due to mechanical stimuli named *osteogenic index (OI)* (Stevens et al., 1999) was calculated for each element within the proximal growth plate. Subsequently, a second FE analysis was conducted to simulate growth of elements taking the element’s growth rate and the direction of growth into account using Abaqus’ (Dassault Systémes Simulia Corp, Rhode Island, USA) orthotropic thermal expansion (Kainz et al., 2020; Yadav et al., 2016). Three different approaches to model growth directions have been proposed in the literature (Carriero et al., 2011; Hunziker, 1994; Yadav et al., 2016). A figure visualizing the three different growth direction methods is included in the supplementary material.

a. Femoral Neck Deflection Direction (FNDD): A uniform growth direction was calculated based on the average neck deflection direction during loading. It was calculated as the mean vector between a node at the neck base and the femoral head center during the nine load instances. This growth direction was applied to all elements in the second FE analysis.
b. Principal Stress Direction (PSD): For each element, the growth direction was defined as the direction of the highest principal stress. Therefore, each element had a unique growth direction vector.
c. Normal to growth plate orientation (NORM): A uniform growth direction was calculated as the normal vector to the main orientation of the growth plate obtained using a principal component analysis. This growth direction was applied to all elements in the second FE analysis.

The AVA was identified by calculating the angle in transverse plane between the neck axis (i.e. vector between nodes representing the femoral neck base and the femoral head center) and the knee axis (i.e. vector between nodes representing the epicondyles). Doing so with the same nodes in the model before and after growth allows to quantify the predicted development of AVA by the mechanobiological multi-scale workflow. A visual overview of both finite element analysis and the quantification of AVA is shown in Figure 3.

**Figure 3:**
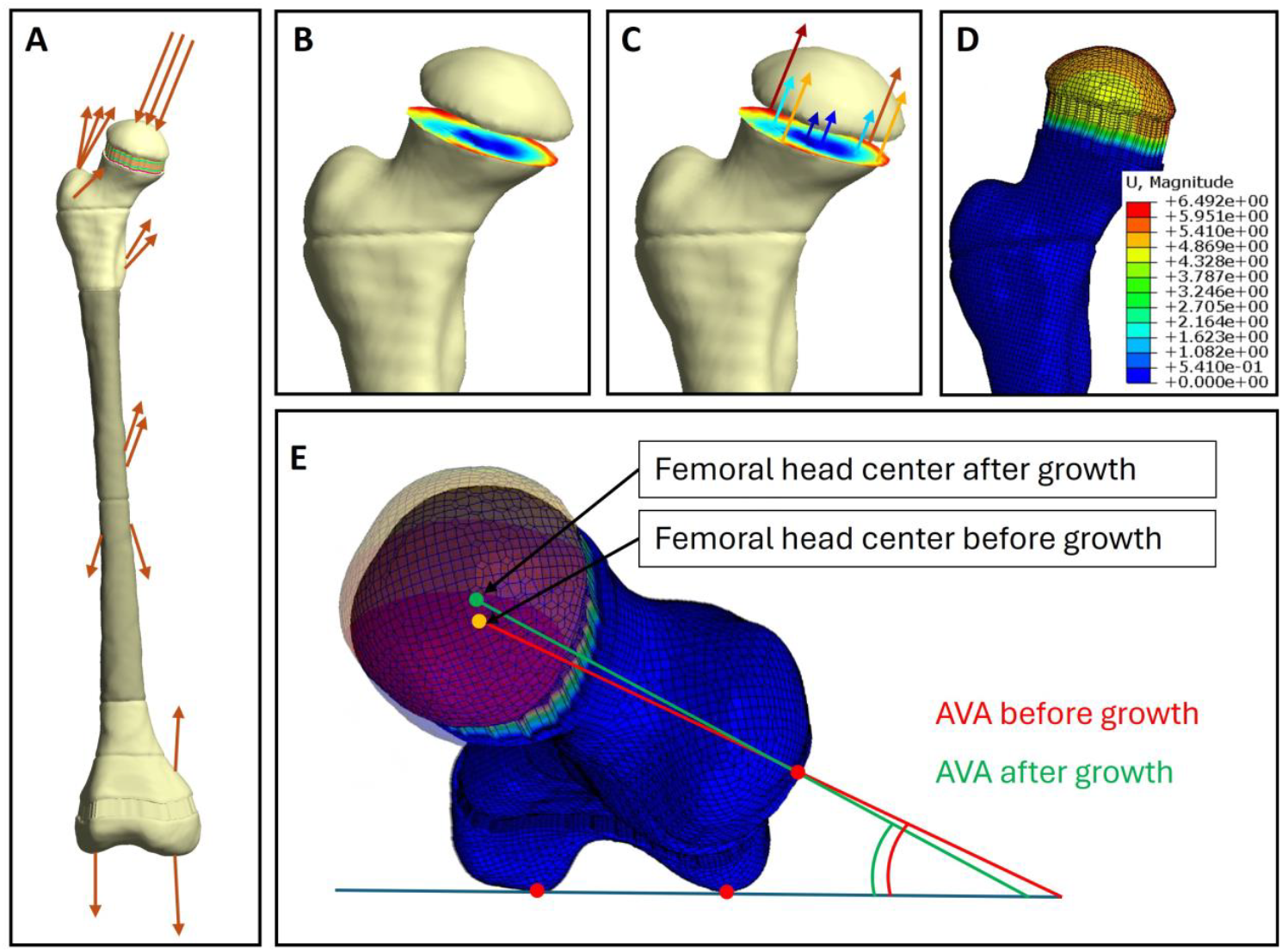
Visual description of the methodology to model growth and quantify the change of anteversion angle (AVA). Subfigure **A** shows the finite element analysis with muscle and hip joint contact forces applied to the femur. Subfigure **B** represents the calculation of the growth rate for each element within the growth plate based on the mechanical shear and compressive stresses and several influencing factors discussed in section 2.5. Subfigure **C** shows a simplified representation of the growth rate applied through orthotropic thermal expansion. The direction in which growth is modeled can vary depending on the chosen methodology (FNDD, PSD, NORM). In subfigure **D** the displacement due to modeled growth is visualized. Subfigure **E** shows the baseline and the grown model overlayed from a top view. The location of femoral head center is moved while other nodes stay at the same location. The change of AVA is calculated as the difference between the AVA before and after growth.

### 2.5. Sensitivity analysis

As the result of the mechanobiological simulation workflow depend on various parameters, a sensitivity analysis was performed to evaluate their influence. In total, 330 multi-scale simulations were performed for each femur and for each of those, the predicted development of AVA was evaluated.

Firstly, two loading conditions (SO and EMG-informed) were derived from MSK simulations for each femur and used to identify shear and hydrostatic stresses within the proximal femoral growth plate.

Secondly, with those stresses the growth rate was calculated for each element *i* within the growth plate using different parameter variations. In general, the growth rate includes a weighted linear combination of shear (*σ*_*S*_) and hydrostatic (*σ*_*H*_) stresses representing the OI (i.e. growth due to mechanical stimuli) (Equation 1) and a value representing the biological growth *growth*_*bio*_ (Equation 2). The ratio of weighting factors *a* to *b* for shear and hydrostatic stresses was varied (0.02, 0.1, 1/7, 0.25, 0.5, 1, 2, 4, 7, 10, 50). Additionally, we introduced parameter sets where negative growth rates were avoided or where the growth rate was normalized to a specified range to account for differences in the magnitude of the OI. For each of the eleven ratios, the following growth simulations were performed:

d) Including biological growth where the amount was twice the maximum mechanical growth as in previous studies, i.e. Equation 2 with *growth*_*bio*_ = 2 * *max*(*OI*)
e) Neglecting biological growth, i.e. Equation 2 with *growth*_*bio*_ = 0
f) Neglecting biological growth but avoiding negative growth, i.e. Equation 3
g) Neglecting biological growth but normalizing growth rate, i.e. Equation 4
h) Including biological growth and normalizing growth rate, i.e. a) and Equation 4

This resulted in 55 unique parameter combinations which led to different growth rates depending on the shear and hydrostatic stresses at the growth plate. Combining each of these 55 parameter variations with each of the loading conditions (SO and EMG-informed) resulted in 110 simulations.

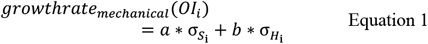

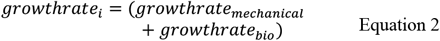

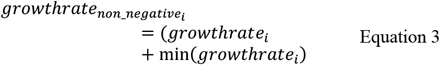

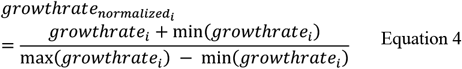

Thirdly, three approaches how the growth direction can be modeled are proposed in literature. Therefore, we decided to run each of the so far 110 simulations with each growth direction method (FNDD, PSD, NORM), which resulted in abovementioned 330 simulations for each femur.

### 2.6. Data analysis

HJCF magnitude and orientation as well as the sum of all muscle forces acting on the femur during the stance phase were compared between the SO and EMG-informed approach using paired t-tests with Statistical Parametric Mapping (Pataky et al., 2013).

To account for inaccuracies in AVA measurements, 100 imputations of the measured development of AVA were created by adding a random value within the measurement uncertainty range identified as the maximum difference between measurements, i.e. ±3.58°. This resulted in an evenly spaced distribution. A stepwise multiple linear regression using a backward elimination procedure for statistically non-significant predictors (p>0.05) was performed using AVA and NSA from S1, age (at S1 & change between S1 and S2), weight (at S1 & change between S1 and S2), height (at S1 & change between S1 and S2), steps per day and hours of sport per week as predictors and the measured development of AVA as independent variable. All predictors were normalized to their z-scores. The linear regression model was calculated for all 100 imputations (same predictors, randomly imputed independent variable) and the mean of all adjusted R^2^ values was used for further evaluations. The regression model without the multi-scale predictions of AVA development was used as a baseline measure (i.e. *baseline model*).

Furthermore, for each of the 330 growth simulations, the same stepwise multiple linear regression was performed with the multi-scale prediction of the development of AVA as an additional predictor to identify the models’ power to explain the development of AVA (i.e. *explaining model*). Again, these regression models were calculated for all 100 imputations. Subsequently, linear regression models which did not include the multi-scale prediction as a predictor or where this predictor was associated with a negative slope were excluded for further analysis. The linear regression models were ranked depending on their adjusted R^2^ across imputations and the best performing models were analyzed in more detail regarding their similarities and differences within the various parameters of the workflow.

Additionally, the same procedure was performed to identify the models’ power to predict the development of AVA (i.e. *predicting model*), solely by the data from S1, steps per day, hours of sport per week and the multi-scale prediction of the mechanobiological model as this would be data that could be used for clinical decision-making.

### 2.7. Qualitative analysis of best explaining linear regression model and selected femurs

For the single best explaining model, the growth rates for each element in the growth plate were projected on the transverse plane according to the elements’ locations and interpolated to a squared grid. A blue to red color scheme was used to visualize growth rates representing minimum to maximum values, respectively. This resulted in heatmaps of equal size for all growth plates revealing distribution of the growth rate and allowing to qualitatively identify differences between femurs experiencing different growth trends (i.e. decrease or increase of AVA). Additionally, the center of growth was calculated, similarly as one would calculate the center of mass of a disc where the growth rate represents the weight at a location.

## 3. Results

The ten children included in this study, aged 9.9±0.9 years at the first data collection session, grew 13.5±3.3 cm and gained 8.2±3.3 kg of body mass during the two years. Participants’ AVA was 34.7±8.9° at the initial assessment and changed between −13.1° and +11.8° between data collection sessions (Table 1). The average daily step count was 9592 (range: 5671 – 15212) and participants reported to perform sports between 5 and 29 hours per week (mean 13.6 ± 7.8 hours).

### 3.1. Muscle and joint contact forces estimated with different approaches

The EMG-informed MSK simulations, where muscle activations of the model had to follow the shape of EMG-recorded muscle activations additionally to effort minimization, led to significant different (p<0.05) muscle forces compared to the SO approach, which minimizes the sum of all muscle activations (Figure 4). The forces produced by musculus tensor fasciae latae, the hamstring muscles and the knee extensor muscles were significantly increased during the complete stance phase in the EMG-informed simulations compared to SO simulations. The soleus produced significantly less force while the gastrocnemius forces were significantly increased in EMG-informed simulations compared to SO simulations. Unfortunately, the EMG data of one participant were not of sufficient quality and therefore all results involving EMG rely on data of 18 femurs. This participant’s data was removed for performing statistical tests between SO and EMG-informed approaches.

**Figure 4:**
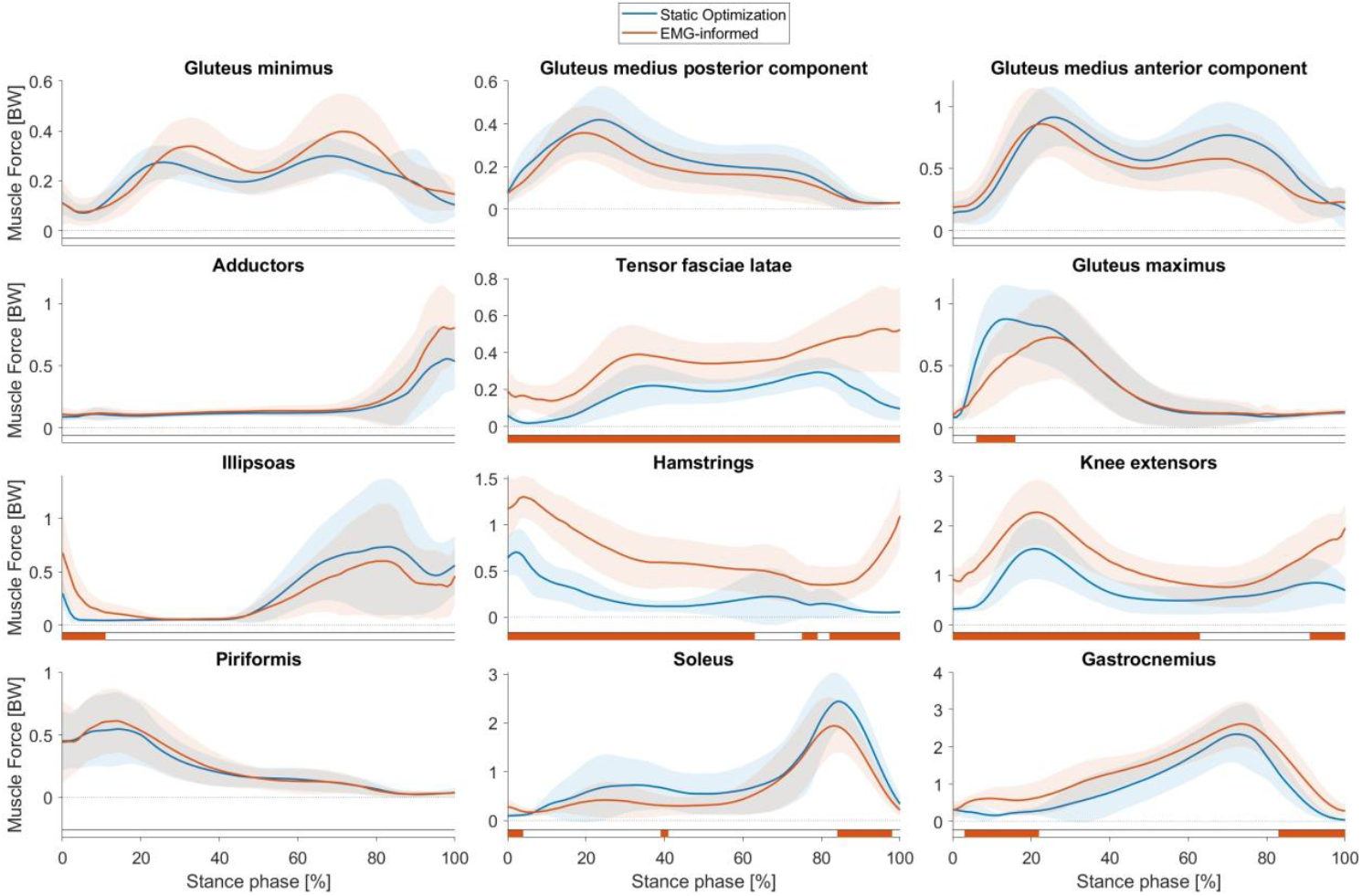
Muscle forces estimated by musculoskeletal simulations with two approaches to solve the muscle redundancy problem, i.e., static optimization (blue waveforms) and EMG-informed (red waveforms) approach. Significant differences (p<0.05) between both approaches identified with Statistical Parametric Mapping are visualized by orange-colored bars beneath each subplot.

The magnitude of hip, knee and patellofemoral joint contact forces were significantly higher in EMG-informed simulations compared to the simulations performed with SO (Figure 5). At the hip, mainly the vertical component was increased, whereas at the knee joint, all components are significantly higher. The orientation of the HJCF was only significantly different between both modelling approaches in small parts of the stance phase (Figure 5).

**Figure 5:**
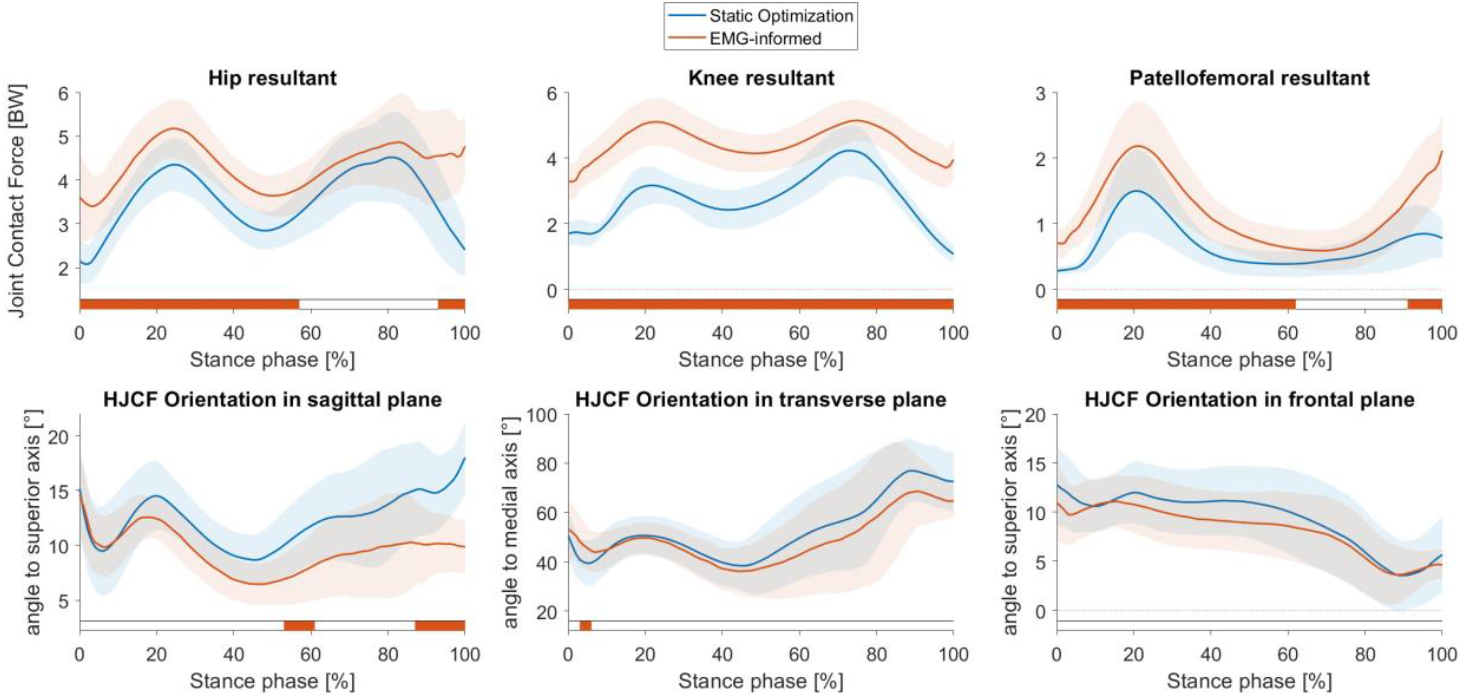
Joint contact forces and hip joint contact force (HJCF) orientation estimated by musculoskeletal simulations with two approaches to solve the muscle redundancy problem, i.e., static optimization (blue waveforms) and EMG-informed (red waveforms) approach. Significant differences (p<0.05) between both approaches identified with Statistical Parametric Mapping are visualized by orange-colored bars beneath each subplot.

### 3.2. Sensitivity analysis with mechanobiological growth predictions

The *baseline* linear regression model, which did not include multi-scale predictions for AVA development, was highly significant (p<0.01) and showed a moderate mean coefficient of determination (R^2^=0.46) across the imputations. The AVA and the age at the first session, as well as change of weight between sessions were significant predictors with a negative slope. Average steps per day and the change of height between sessions were significant predictors with a positive slope.

In 140 linear regression models, the predicted development of AVA from the mechanobiological multi-scale workflow was excluded as a significant predictor during the stepwise backward elimination procedure, therefore these models were equal to the *baseline* model. One linear regression model included the multi-scale prediction with a negative slope, meaning that the predicted and measured change of AVA was inversely correlated, and was therefore excluded from further analysis. This model’s adjusted R^2^ across imputations was 0.65, therefore, not within the 10 best explaining models. 189 analyses remained for further examination. The 10 best performing models, i.e. models with the highest adjusted R^2^ across imputations (R^2^>0.685), used muscle forces and HJCF estimated with SO as loading condition to calculate stresses within the growth plate. In all of those models, negative values as growth rate were allowed. Furthermore, seven of the 10 best performing models did not account for biological growth and in six, the growth rate was normalized to a specific range. Regarding the ratio of weighting factors *a* to *b* for shear and hydrostatic stresses, respectively, no clear trend was observed. In eight of the 10 best performing models, the NORM growth direction method, in which the direction of growth is modelled normal to the orientation of the growth plate, was used (Figure 6). A figure visualizing the distribution of the predicted AVA development from all 330 parameter sets is included the supplementary material.

**Figure 6:**
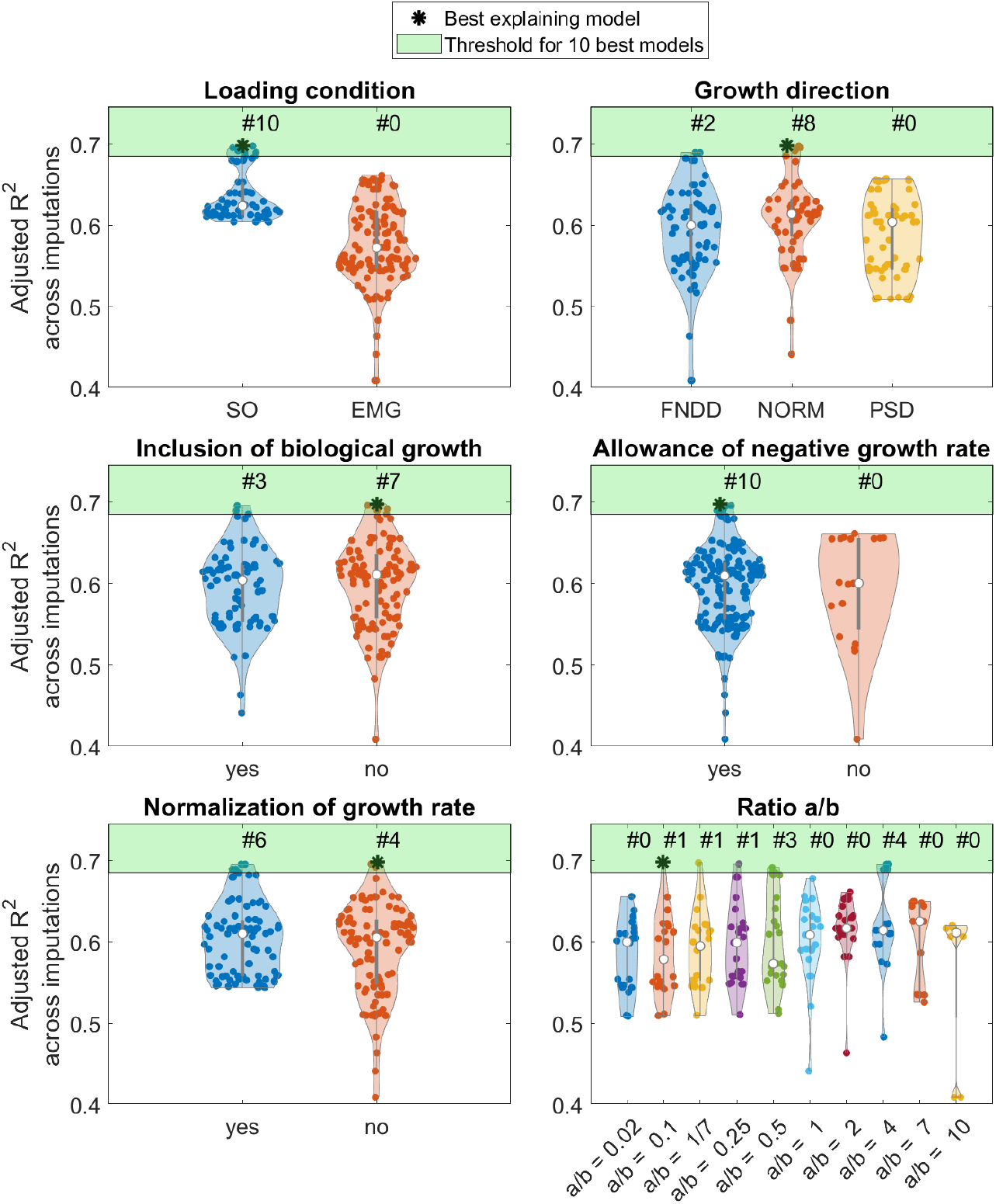
Adjusted R^2^ of the linear regression models and parameters used for mechanobiological multi-scale simulations. The light green patch highlights the area above the threshold for the 10 best explaining models.

### 3.3. Analysis of best explaining and predicting model

The model with best explanation power was a refinement of the *baseline* model with the multi-scale prediction and the NSA at the initial assessment as additional significant predictors with positive and negative slopes, respectively. The loading obtained from MSK simulations with SO were used as input to estimate the stresses within the growth plate. Biological growth was not included, values were not normalized to a specified range and the growth rate values were allowed to be negative. The ratio of weighting factors *a* to *b* was 0.1. The multiple linear regression model was highly significant (p<0.001) and showed a high coefficient of determination across imputations (R^2^=0.7). In 18 femurs, the development of AVA was explained successfully in terms of direction. In the remaining two femurs, the measured development was low with values of −2.1° and +1.7° (Figure 7 left).

**Figure 7:**
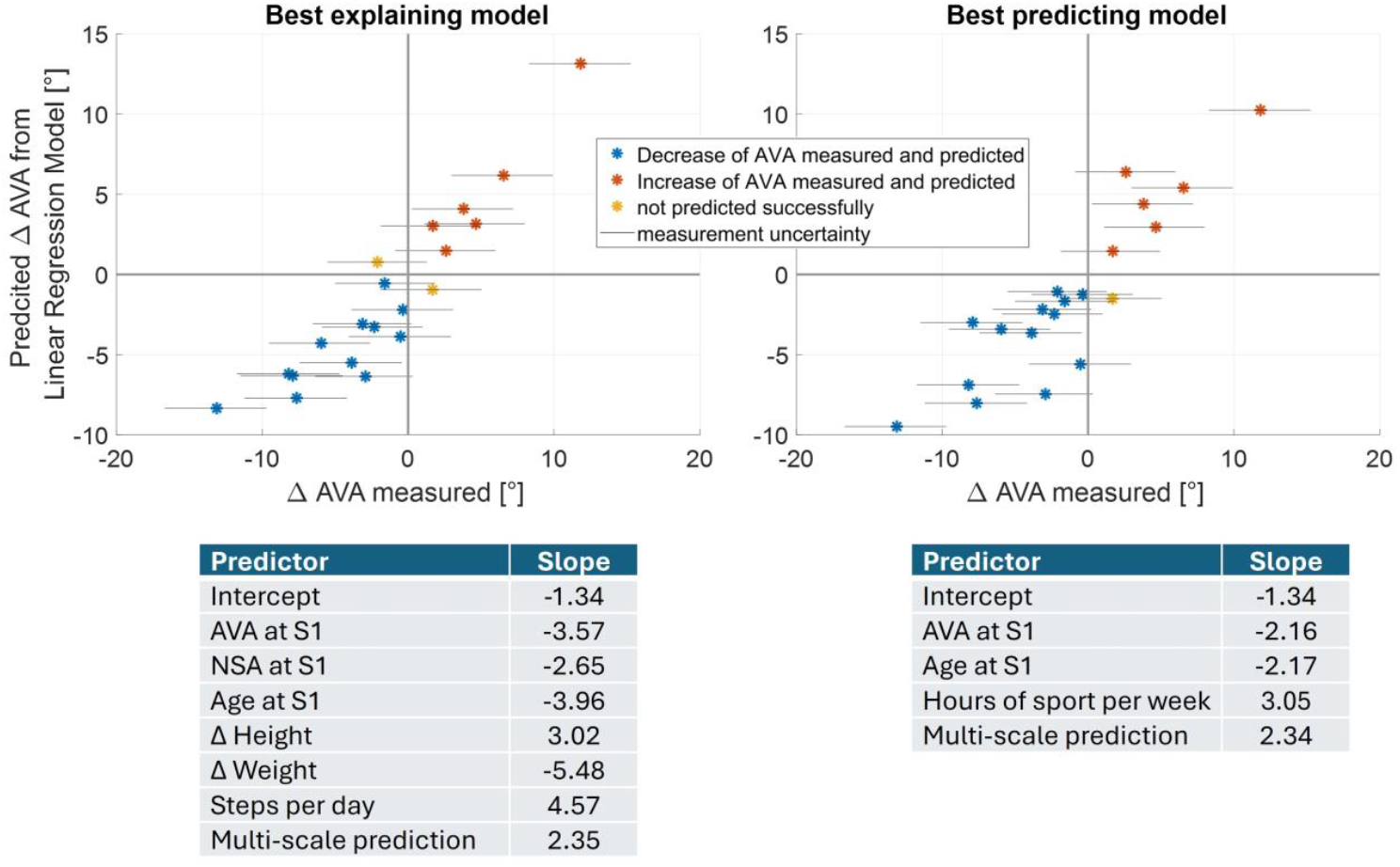
Scatter plot based on the measured anteversion angle (AVA) and predicted development of AVA for best explaining (left) and best predicting (right) model. The stars indicate the measured change of AVA assessed under the two-person verification principle and the horizontal lines passing the colored markups indicate measurement uncertainties. The tables below include the significant z-normalized predictors and their slopes of the corresponding model. S1 = first data collection session.

The best performing predicting model (i.e. based solely on data from the initial data collection and multi-scale predictions) was highly significant (p<0.001) with almost as high coefficient of determination (R^2^=0.66) across imputations as the best explaining model (Figure 7 right). It included only the age and the AVA from the initial assessment as significant predictors with negative slopes, as well as the hours of sport per week and the multi-scale predictions as significant predictors with a positive slope. The parameters of the multi-scale workflow were similar to those of the best explaining model expect for the ratio of weighting factors *a* to *b*, which was 0.02.

### 3.4. Qualitative analysis of best explaining model

Heatmaps visualizing the growth rates of all femurs showed no clear trend and indication whether an increase or decrease was predicted by the multi-scale simulations (Figure 8). A significant correlation (p<0.05) was found between center of growth in the medial/lateral direction and the predicted development of AVA with a low coefficient of determination (R^2^=0.25). No significant correlation was found between the prediction and the center of growth rate in anterior/posterior direction.

**Figure 8:**
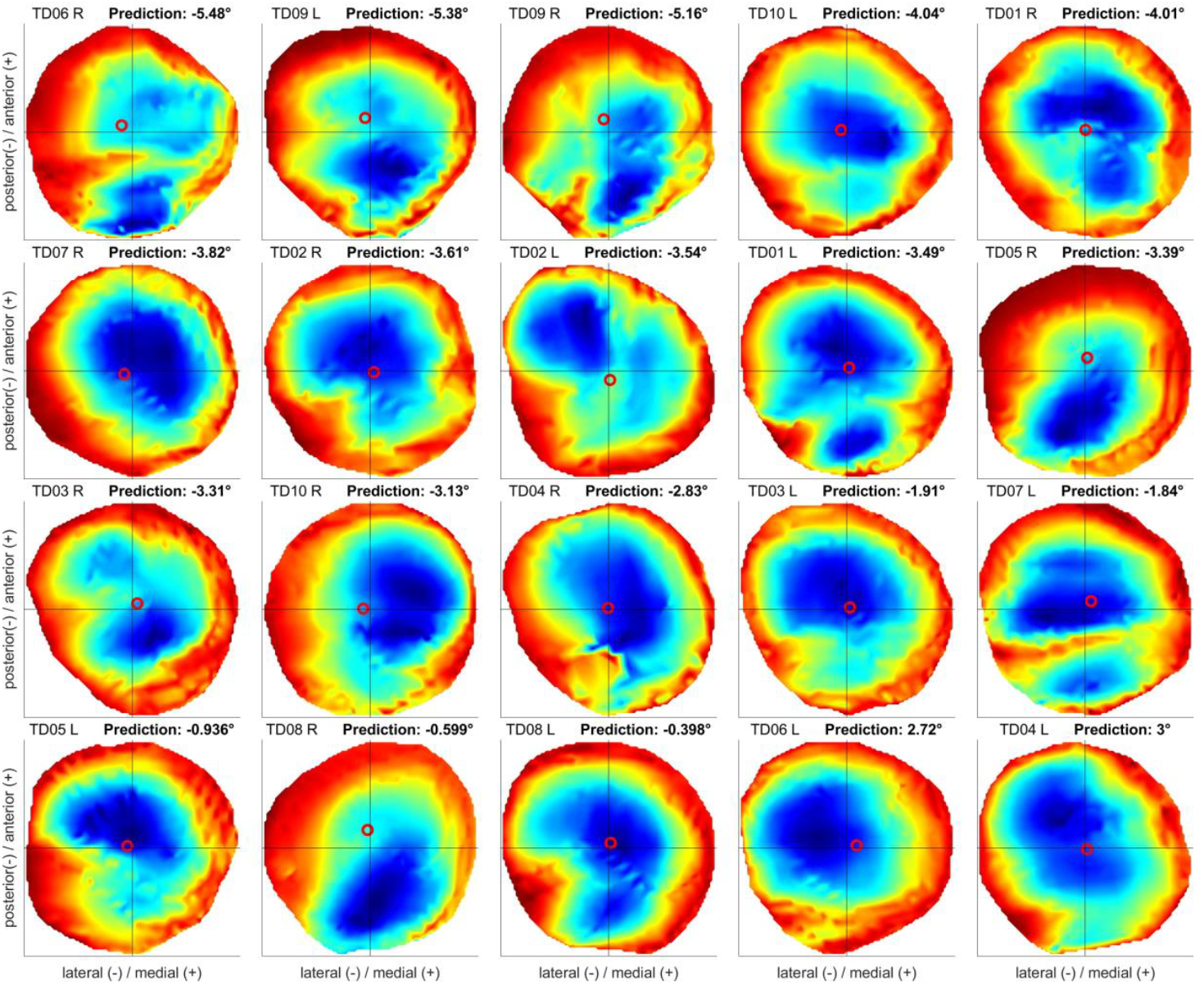
Growth rates of the best explaining model within the proximal femoral growth plate visualized using a blue to red color scheme representing low and high values, respectively. The red circle is indicating the center of growth rate. The order of the diagrams is based on the predicted development of AVA by the multi-scale simulation (left to right, line by line). A figure with a unique color scheme across all heatmaps is included in the supplementary material.

## 4. Discussion

We experimentally investigate a mechanobiological model to predict the development of femoral AVA with the use of MRIs captured at two occasions approximately two years apart. The sensitivity analysis identified best parameter combinations for the mechanobiological model. Linear regression models were able to explain (R^2^=0.7) and predict (R^2^=0.66) the development of the AVA based on the mechanobiological simulations with high coefficient of determinations.

The inclusion of the mechanobiological multi-scale predictions increased the explanatory power of the linear regression model compared to the *baseline* model. 70% of the development of AVA were explained by the best model including physical activity and anthropometric data of the first session and the changes between sessions as well as the multi-scale predictions. A model including only data of the first session and the multi-scale predictions had only 4% lower explanatory power than the best explaining model. Considering that bone growth is guided by mechanical loading (Carter et al., 1996; Carter and Beaupré, 2000; Wolff, 1892), but keeping in mind that there are other factors like nutrition or genetics which influence biological tissue responses (Stevens et al., 1999), the observed coefficient of determinations can be considered as high.

The shape and magnitude of muscle forces obtained with personalized musculoskeletal models and SO as approach to solve the muscle redundancy problem, were in agreement with previous studies (Lin et al., 2012; Trinler et al., 2019). The two approaches (SO and EMG-informed) to solve the muscle redundancy problem in musculoskeletal simulations estimated significantly different muscle and joint contact forces. The use of EMG data to inform simulations increased mainly muscle forces of the tensor fasciae latae, hamstring and knee extensor muscles.

This has also been observed in other studies and is possibly due to a higher co-contraction in some participants (Hoang et al., 2019). As a consequence of higher muscle forces spanning the hip and knee joints in the EMG-informed simulations, the magnitude of hip, knee and patellofemoral joint contact forces were significantly higher in EMG-informed simulations compared to those obtained with the SO approach. Furthermore, the orientation of the HJCF in the sagittal plane differed between both modelling approaches. As a consequence, different nodes of the femoral head are loaded which in turn modifies the stress distribution.

The sensitivity analysis revealed that multi-scale predictions based on the mechanical loading estimated with SO show higher coefficients of determination than those that used muscle and joint contact forces obtained with an EMG-informed approach. This is in contrast to our expectations because studies have shown that EMG-informed simulations increase the agreement between simulated and in-vivo measured joint loadings due to the consideration of subject-specific motor control (Bennett et al., 2022; Hoang et al., 2019; Manal and Buchanan, 2013). However, in our study we analyzed healthy children with typical walking patterns. Walking in humans is a very efficient task and therefore low co-contraction can be assumed (Anderson and Pandy, 2001; Waters et al., 1988). Comparing muscle activation patterns obtained with SO with experimentally measured EMG signals showed good agreement in our participants. Therefore, the results obtained using SO serve as a valid estimation for the investigated cohort. It might be that EMG-informed simulations better estimate the loadings in pathological populations, e.g. in participants with neurological disorders like cerebral palsy (Wesseling et al., 2020).

Growth simulated in the NORM direction led to the most accurate predictions. In most models with high explanatory power, the growth was simulated in NORM followed by the FNDD direction. As the growth direction of the FNDD method is highly influenced by mechanical properties of the FE model, we would encourage peers to use the NORM method for future studies because it demonstrates superior performance with reduced uncertainty compared to the other growth direction methods. Furthermore, it seems that the orientation of the growth plate has a big impact on the growth direction of the proximal femur. Nevertheless, it needs to be investigated if the NORM method also performs best in pathological cases in future studies.

The mechanobiological model includes many parameter combinations, which were analyzed with our sensitivity analysis simulations. The best explanatory and predicting models had a ratio of weighting factors *a* to *b* of 0.1 and 0.02, respectively, indicating that the factor *b* for the hydrostatic stress should be higher than *a* to ensure best predictions. While this is in contrast to the ratios used in previous cross-sectional studies using the multi-scale mechanobiological workflow (Carriero et al., 2011; Kainz et al., 2020; Koller et al., 2024, 2023b; Shefelbine and Carter, 2004; Yadav et al., 2021, 2017, 2016) the visualization of the growth rates as heatmaps showed similar patterns compared to the previous investigations, i.e. ring-shaped with high values on the outside and low values at the center, and therefore might not impact the conclusions of previous studies. The inclusion of a generic value accounting for the biological growth, the normalization of growth rates to a specified range as well as the restriction to positive growth rate values did not increase the accuracy of the predictions.

The visualized growth rates of the best explaining model did not reveal a clear distinction in terms of shape or distribution between individuals experiencing an increase or a decrease of AVA (Figure 8). The center of growth rate in these heatmaps could indicate to which direction the femoral head will tilt in the growth simulations. However, a significant but only low correlation was found between the weighted center of growth rate in medial/lateral direction and the predicted development of AVA. In detail, a more medially pronounced growth rate distribution correlated with an increase of AVA.

During skeletal growth, the AVA typically decreases from 40° to approximately 20° (Bobroff et al., 1999; Fabry et al., 1973). We analyzed longitudinal data of healthy children and found an increase of AVA between sessions in some femurs, which is considered as unphysiological growth from a clinical point of view. The detailed analysis of femurs that experience pathological increase of AVA in terms of growth plate morphology and loading conditions did not reveal key factors that might explain the reason for this pathological development. This strengthens the need for the performed multi-scale simulations as it includes the subject-specific overall femoral geometry, the shape and orientation of the growth plate and the mechanical loading on the growth plate during walking to simulate the cells biological response to these stresses. Only accounting for all these aspects collectively led to the observed high coefficients of determination.

The used multi-scale mechanobiological model includes several simplifications and did not aim to mimic real femoral development, which includes surface bone re-modelling additionally to growth at the growth plates. In the used model, growth was only simulated at the proximal growth plate while all other elements were kept constant. Therefore, all geometrical changes result from a tilt and shift of the femoral head proximal to the growth plate, e.g. an increase of AVA would be represented by an anterior tilt of the femoral head. While this modelling choice did not include any surface bone re-modelling and adaptations, the model was sophisticated enough to fulfil its purpose to identify femoral AVA growth trends. Furthermore, previous studies showed that this model can distinguish between healthy and pathological femoral development (Carriero et al., 2011; Kainz et al., 2021; Koller et al., 2024, 2023b). Due to the limitation of modelling growth only at the proximal growth plate, we decided to compare only the clinical important feature, i.e., AVA, between the measured and predicted values instead of comparing the entire shape of the bone. In the future, we plan to develop more realistic growth models that predict the full development of the femoral geometry. Methods such as Generalized Procrustes Analyses (Bastir et al., 2024) can then be used to compare the predicted with the actual femoral geometry.

Negative growth and ignoring biological growth is certainly not physiological plausible. Negative growth rates are theoretically possible because hydrostatic compressive stress is negative. Hence, if the negative term 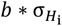 is absolutely speaking larger than the term 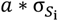 the resulting growth rate for this element is negative. The model is based on experimental observation which showed that growth plate cartilage ‘likes’ compressive stress and therefore does not lead to ossifications, whereas the cartilage does not ‘like’ shear stress and therefore leads to ossification (Stevens et al., 1999). Therefore, negative growth rates indicate that growth is not promoted due to mechanical stimuli at these elements. However, if biological growth would be considered, the total growth would still be positive. Similar to previous studies, biological growth was modeled as a simple constant value. Surprisingly, ignoring the biological growth rate and allowing negative growth rates led to the best results, indicating that modelling the biological aspect with a simple constant value does not work very well and leads to worse predictions. Hence, future work is needed to come up with better and potentially subject-specific ways to include the biological growth amount.

This study included several limitations. Firstly, only data of TD children were included in this study. Collecting longitudinal data of participants with bony pathologies (e.g., cerebral palsy, skeletal dysplasia, rickets, etc.) is challenging because these patients typically undergo diverse clinical interventions, including de-rotation osteotomies, within such a long period of time. Nevertheless, a wide range of AVA changes were observed within the investigated cohort, which included typical but also pathological AVA developments. Secondly, the investigated cohort was relatively old considering that changes of AVA are higher in early childhood. Nevertheless, several participants experienced a growth spurt accompanied with substantial development of AVA within the time period between data collection sessions. However, the identified best parameters might differ for other age groups. Thirdly, similar to previous studies (Kainz et al., 2020; Koller et al., 2024, 2023b; Yadav et al., 2021, 2017, 2016) generic linear-elastic material properties were used for all parts of the femur in the FE model. Fourthly, the model did not include all anatomical details, e.g., the Ring of Lacroix around the growth plate was not included. Previous studies, however, showed that the Ring of Lacroix has only a minor impact on the stress distribution (Hucke et al., 2023; Piszczatowski, 2012). Hence, we kept the model similar to previous investigations, which already revealed plausible simulation results (Kainz et al., 2020; Koller et al., 2024, 2023b; Yadav et al., 2021, 2017, 2016). Fifthly, we used boundary conditions similar to previous studies (Kainz et al., 2020; Koller et al., 2024, 2023b; Yadav et al., 2021, 2017, 2016). However, a recent study investigated different boundary conditions in FE analysis of the femur and proposed a new method to represent the biomechanical loading situation more accurately (Bavil et al., 2024). Unfortunately, this new boundary condition restricts movement of selected nodes close to the proximal growth plate and therefore induces artificial stresses within the growth plate. Therefore, it was not suitable for this study’s modelling purposes. Importantly, the GP-Tool, which we developed and used in this study to create the FE models is open source and different boundary conditions can be implemented easily and used in future studies.

In conclusion, the findings of this study showed that multi-scale simulations combined with a mechanobiological model enables an accurate prediction of femoral AVA growth trends in children. No key factors were identified that could differentiate between healthy and pathological growth patterns, thereby reinforcing the necessity for the multi-scale workflow, which considers multiple factors collectively. The application of mechanobiological model-based predictions may facilitate a paradigm shift in the clinical management of femoral torsional deformities. In contrast to current reactive, invasive procedures such as de-rotation osteotomy, the prospective identification of abnormal loads (which lead to bony deformities) at an early stage could facilitate the implementation of proactive, non-invasive interventions. In the event of pathological bone predictions, the loadings on the bones could be modified through non-invasive interventions, such as gait retraining (Kainz et al., 2024b; Uhlrich et al., 2022), to ensure that the bone loads and growth are within normal range (Kainz et al., 2024a). However, to achieve accurate subject-specific predictions in the future, it is essential to investigate how certain model parameters influence simulation results. This study is the first step towards this goal but further studies including larger sample sizes and pathological cohorts are required to overcome several hurdles before this approach can be adopted in clinical practice. It is essential to identify the range of loading conditions that promote normal bone growth and to determine how participants can achieve these conditions.

