## Supplementary Material for "Femoral bone growth predictions based on personalized multi-scale simulations: Validation and sensitivity analysis of a mechanobiological model"

### 1 Differences between measurements of AVA

For each femur, the points were selected by one researcher with four years of experience in bone segmentation and anatomical feature quantification. The selected points were reviewed following the two-person verification principle by the initial researcher and a second researcher with one year of experience in bone segmentation and anatomical feature quantification. In case of disagreement, points were repositioned with consent of both researchers. The second measurement was used for further analysis and the maximum difference of the change of AVA between the two measurements was used as a measurement uncertainty (Figure S1).

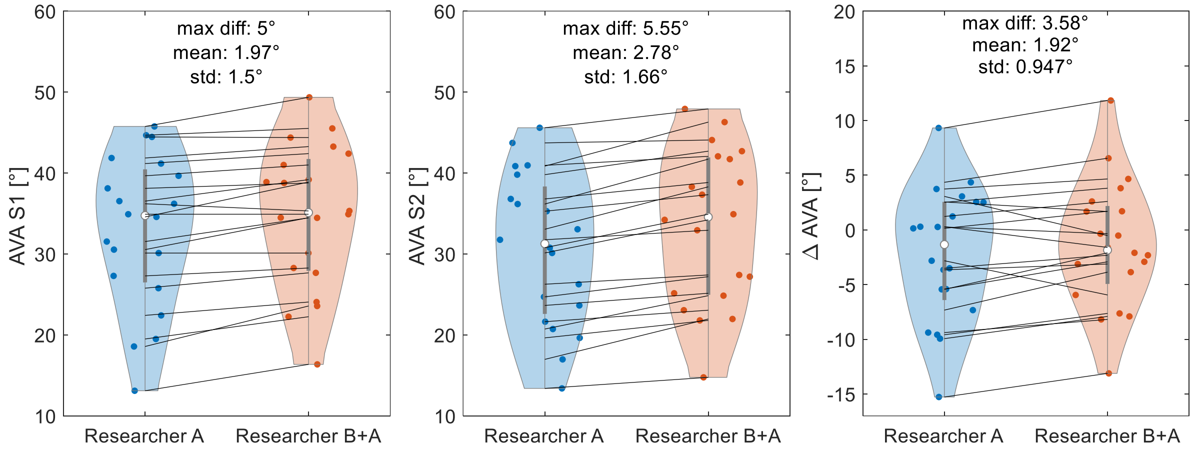

*Figure S1: Differences of femoral anteversion angle (AVA) between both measurements. For each femur, the points were selected by one experienced researcher (A). The selected points were reviewed following the two-person verification principle by the initial researcher and a second researcher (B). In case of disagreement, points were repositioned with consent of both researchers. The second measurement was used for further analysis and the maximum difference of the change of AVA between the two measurements was used as a measurement uncertainty. S1 = first data collection session. S2 = second data collection session.*

### 2 Visualization of different growth direction modeling approaches

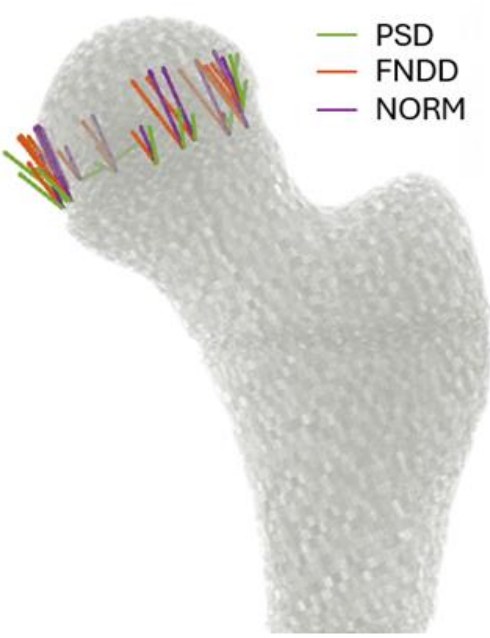

Figure S2 shows the growth rate and direction applied to a subset of elements in the growth plate depending on the used growth direction method. With Principal Stress Direction (PSD), each element has its unique growth direction. With Femoral Neck Deflection Direction (FNDD) and normal to the growth plate (NORM), all elements have identical growth directions.

*Figure S2: The applied growth rate and direction for a subset of the elements in the growth plate are visualized across all three growth direction modeling approaches. With Principal Stress Direction (PSD), each element has a unique growth direction defined as the direction of the highest principal stress. With Femoral Neck Deflection Direction (FNDD), a uniform growth direction is calculated based on the average neck deflection direction during loading. With Normal to growth plate orientation (NORM), a uniform growth direction is calculated as the normal vector to the main orientation of the growth plate obtained using a principal component analysis. Importantly, the amount of growth for each element was based on the osteogenic index calculation, which was independent of the growth direction model.*

#### 3 Detailed analysis of femurs experiencing increase or decrease of anteversion angle

Due to the fact that an increase of anteversion angle (AVA) with skeletal growth is clearly a pathological development [1,2], those femurs that experienced an increase of AVA, even if accounted for measurement uncertainty, were selected and analyzed in detail. The orientation of the proximal growth plate geometry in respect to the femoral coordinate system and to the orientation of the mean and maximum hip joint contact force was compared between the selected femurs experiencing an increase of AVA, those femurs experiencing a decrease of AVA and those femurs experiencing almost no development. The muscle and joint contact forces were compared between the selected femurs with an increase of AVA and all other femurs.

No clear differences were observed between those individuals with pathological AVA development and normal development in all these measures (Figure S3 - Figure S5).

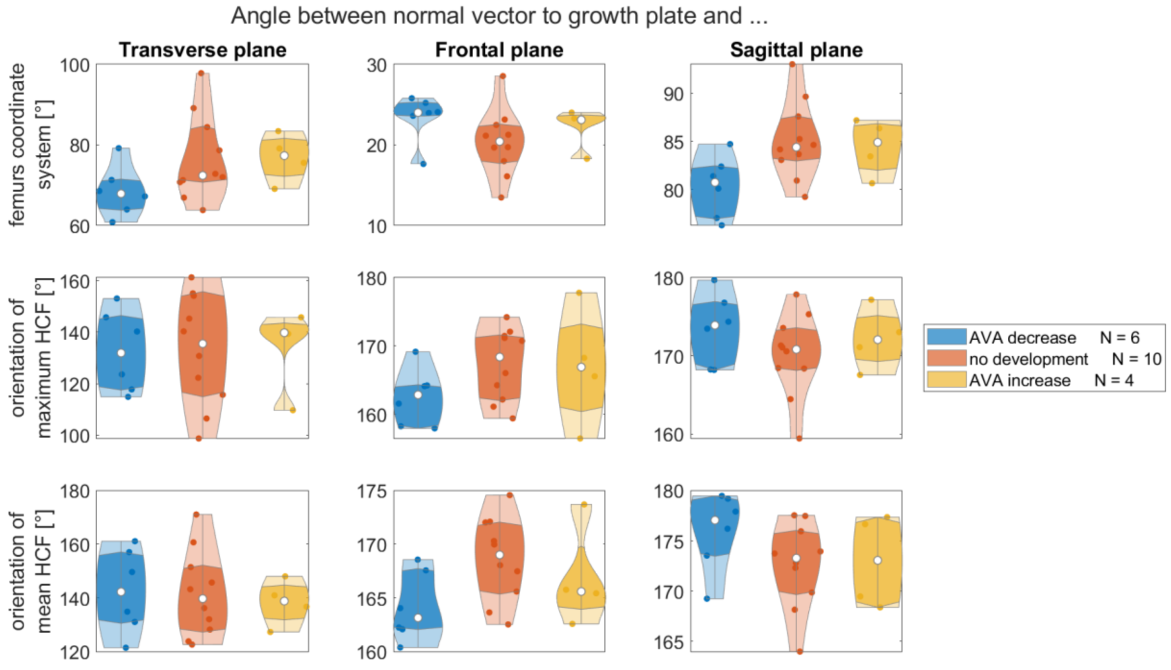

*Figure S3: Comparison of the growth plate orientation (normal vector) between the participants experiencing anteversion angle (AVA) decrease, no development or AVA increase, in relation to the femur's coordinate system (top), the maximum HCF (middle) and the mean HCF during stance phase (bottom) in transverse, frontal and sagittal plane.*

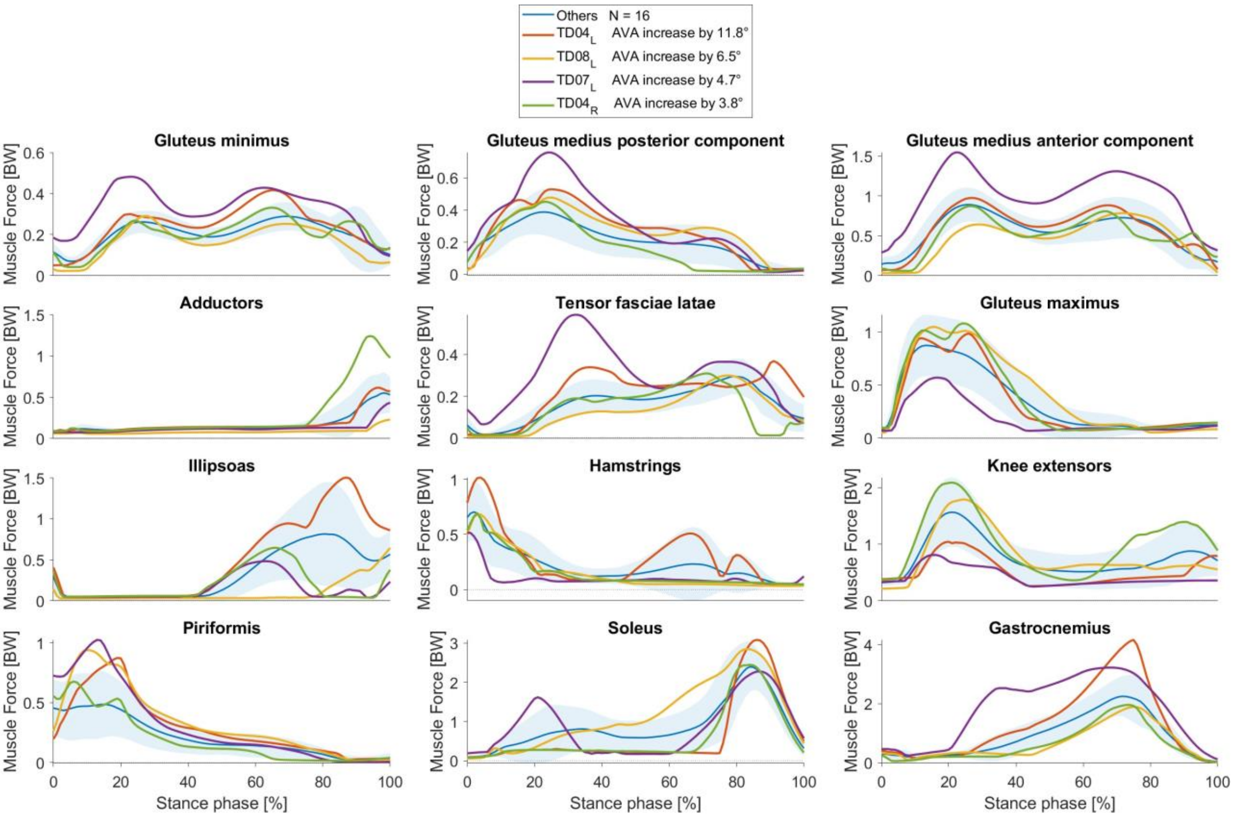

*Figure S4: Muscle forces of individuals experiencing an increase of anteversion angle (AVA) and all others (blue) estimated by musculoskeletal simulations with static optimization to solve the muscle redundancy problem.*

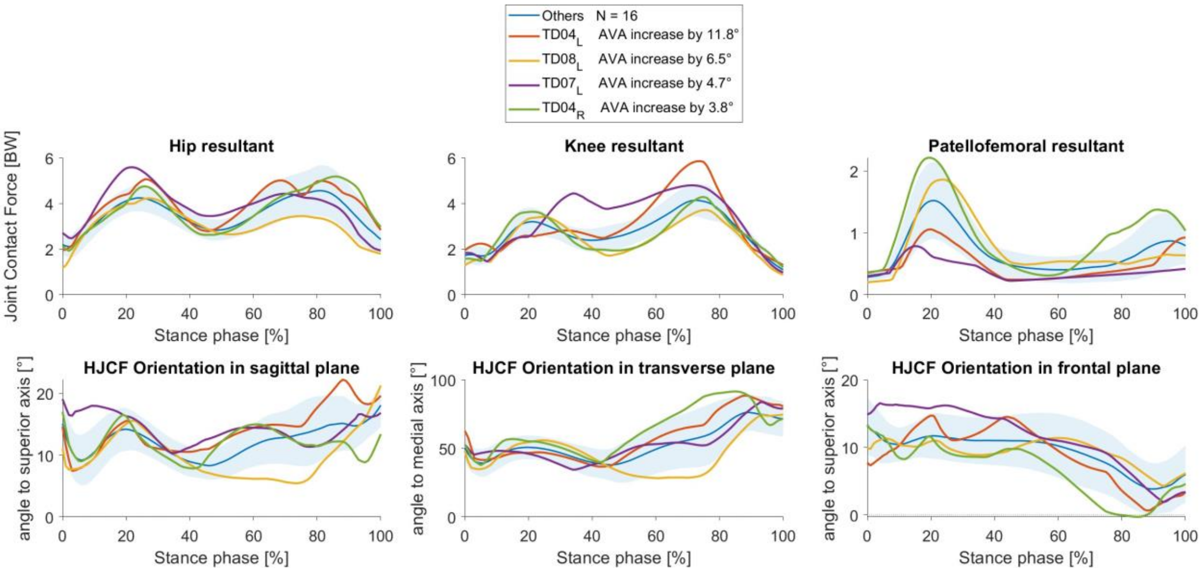

*Figure S5: Joint contact forces of individuals experiencing an increase of anteversion angle (AVA) and all others (blue) estimated by musculoskeletal simulations with static optimization to solve the muscle redundancy problem.*

#### 4 Qualitative analysis of best explaining model

Figure S6 visualizes the growth rates of all femurs as heatmaps similar to Figure 8 of the main manuscript but with a uniform color scheme across all heatmaps.

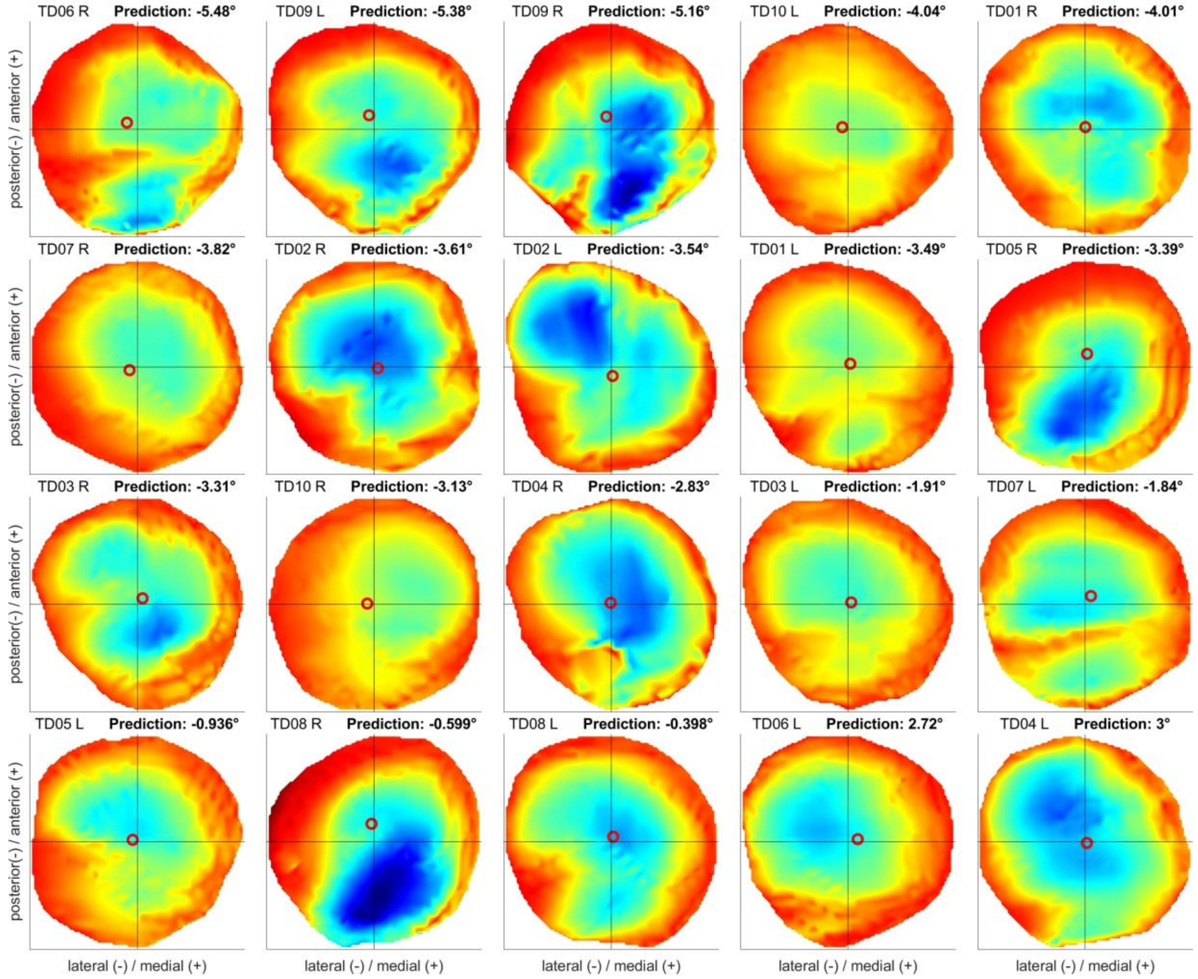

*Figure S6: Growth rates of the best explaining model within the proximal femoral growth plate visualized using a uniform blue to red color scheme across all heatmaps representing low and high values, respectively. The red circle is indicating the center of growth rate. The order of the diagrams is based on the predicted development of AVA by the multi-scale simulation (left to right, line by line).*

#### 5 Distribution of predictions of multi-scale simulations

Figure S7 visualizes the distribution of the predicted development of AVA by the multi-scale simulations with all 330 parameter sets. It shows that in all but one femur, both negative and positive predictions occur. The range of predictions is similarly spread for most femurs, but some participants show a tight range (i.e. TD08_R, left bottom corner) compared to others.

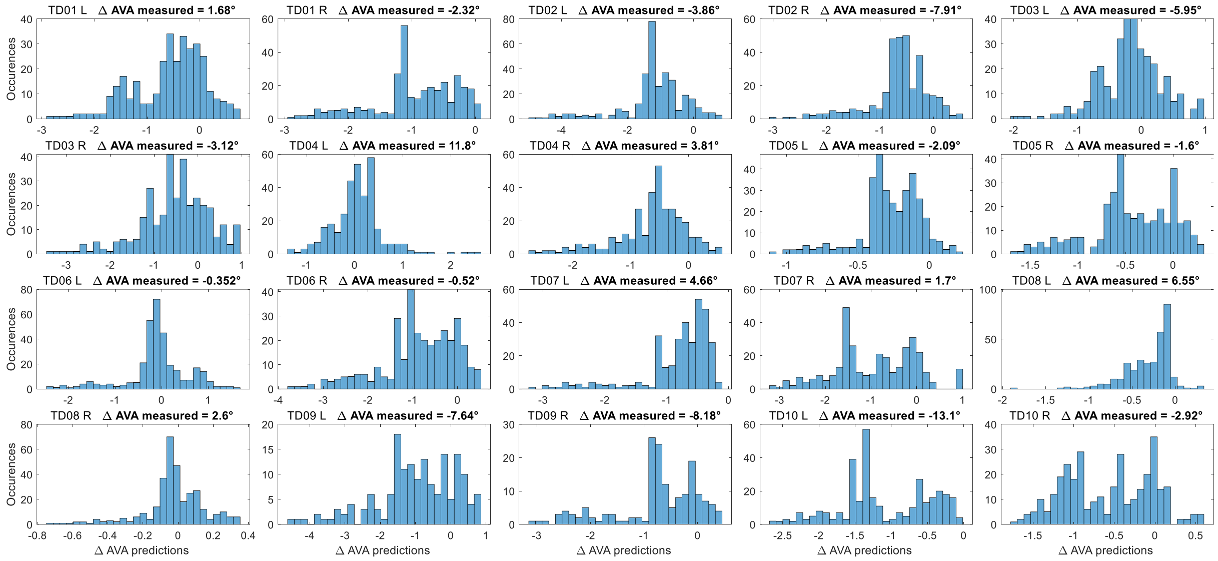

*Figure S7: Distribution of predicted development of AVA by the multi-scale simulations with all 330 parameter sets for each participant.*
